## Supplemental Information for "Human Protein Synthesis Requires aminoacyl-tRNA Pivoting During Proofreading"

**SUPPLEMENTARY METHODS**

All simulations using potential 2 (single Gaussian potential) were performed using the potential V_2_ defined by equation S1:

$$V_{2}=\sum_{bonds} \frac{\varepsilon_{r}}{2}{(r_{i}-r_{i,o})}^{2}+\sum_{angles} \frac{\varepsilon_{\theta}}{2}{(\theta_{i}-\theta_{i,o})}^{2}$$

$$+\sum_{impropers} \frac{\varepsilon_{\chi i}}{2}{(\chi_{i}-\chi_{i,o})}^{2}+\sum_{planar} \frac{\varepsilon_{\chi p}}{2}{(\chi_{i}-\chi_{i,o})}^{2}$$

$$+\sum_{backbone} \varepsilon_{BB}F_{D}(\phi_{i}-\phi_{i,o})+\sum_{sidechains} \varepsilon_{SC}F_{D}(\phi_{i}-\phi_{i,o})$$

$$+\sum_{contacts} \epsilon_{C}C_{W}\left( r_{i,j},r_{i,j,0} \right)+\sum_{non-contacts} \varepsilon_{NC}\left( \frac{\sigma_{NC}}{r_{ij}} \right)^{12}$$

S1

where,

$${\varepsilon F}_{D}\left( \phi\right)=\varepsilon\left( 1-cos\phi\right)+\frac{\varepsilon}{2}\left( 1-cos 3\phi\right)$$

S2

$$C_{W}\left( r_{i,j},r_{i,j,0} \right)=\left( 1+\left( \frac{\sigma_{NC}}{r_{ij}} \right)^{12} \right)\left( 1+W\left( r_{i,j},r_{i,j,0} \right) \right)-1$$

S3

and

$$W\left( r_{i,j},r_{i,j,0} \right)=-exp\left[ \frac{-\left( r_{ij}-r_{ij,0} \right)^{2}}{{2\sigma}^{2}} \right]$$

S4

In these simulations σ is the width of the gaussian well set to a depth of -1, the excluded volume size is σ_NC_ = 2.5 Å, r_ij_ is the distance between atoms I and j, and r_0_ are these distance in the A/A configuration. We performed 100 potential 2 simulations for *H. sapiens* accommodation. In these simulations we reweighted the aa-tRNA contacts with the mRNA by 0.8 and the contacts between the tRNA and ribosome were reweighted by 0.4. These reweighting’s ensure base-pairing between the tRNA and mRNA in these simulations while allowing for reversible fluctuations of the aa-tRNA during accommodation to be consistent with smFRET^1,2^.

Although all structure-based simulations can not capture the chemical reaction of GTP hydrolysis, we started the simulations in the post-GTP hydrolysis state. Therefore, the simulations are starting from a GA state where GTP hydrolysis has already occurred.

**Estimation of aa-tRNA accommodation barrier-crossing activation energy**

The difference in free energy between the A/T aa-tRNA position and the transition state ensemble can be estimated using equation S5:

$k=Ae^{\frac{-E_{A}}{k_{B}T}}$	S5

where, k is the measured rate of the reaction, E_A_ is the activation energy required for the transition from A/T to the transition state ensemble, k_B_ is the Boltzmann constant, and T is the temperature. This equation was used to estimate the E_A_ of human and *E. coli* accommodation. The barrier-crossing attempt frequency (A) was determined as the amount of time required for an accommodation event to occur in simulations, as previously described^3^.

**SUPPLEMENTARY FIGURES**

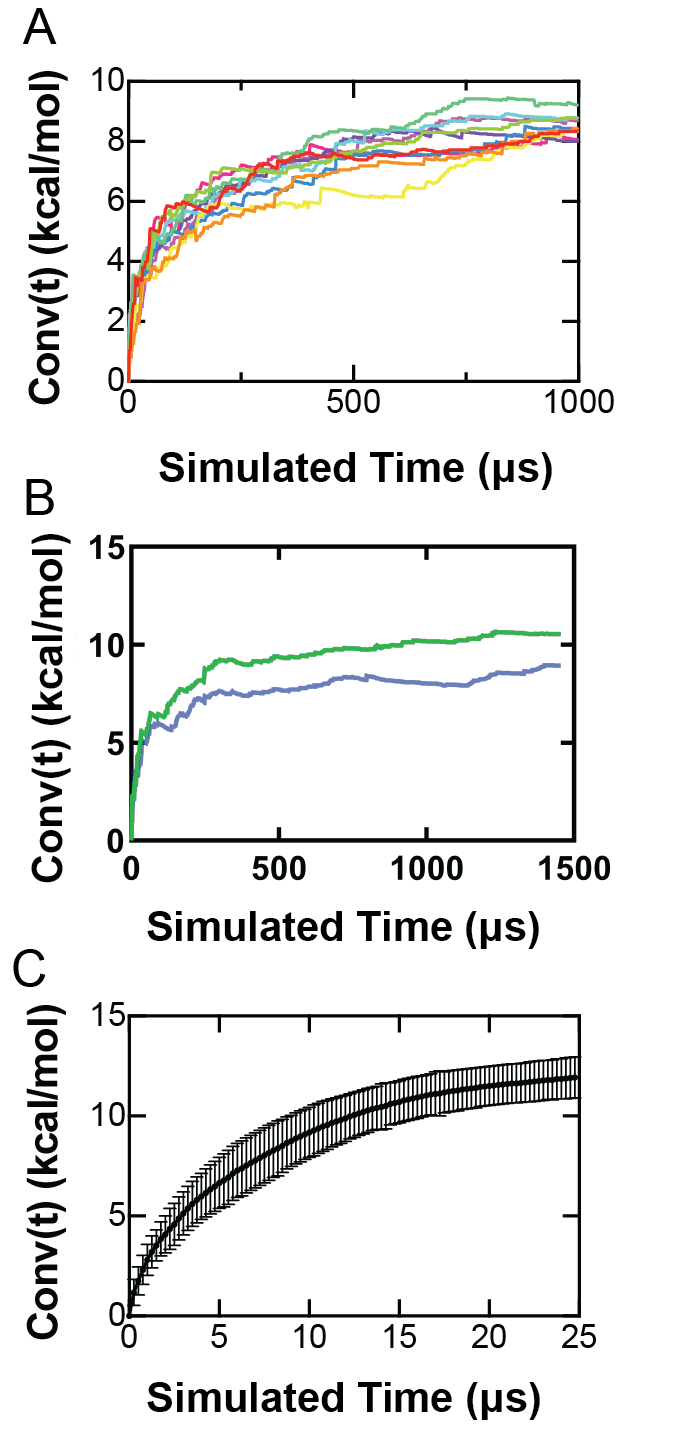

**Supplementary Figure 1 Convergence of structure-based simulations**. (A) Convergence of structure-based simulations as measured by the pointwise RMSD of the free energy landscapes of R_elbow_ and θ_tRNA_ (Conv(t)) for simulations where native contacts are defined using potential 1. (B) Convergence of structure-based simulations as measured by the pointwise RMSD of the free energy landscapes of R_elbow_ and R_CCA_ (blue) or R_elbow_ and θ_tRNA_ (green) for 1.5 ms simulations where native contacts are defined using potential 1. (C) Convergence of structure-based simulations as measured by the pointwise RMSD of the free energy landscapes of R_elbow_ and θ_tRNA_ (Conv(t)) for simulations where native contacts are defined by potential 2.

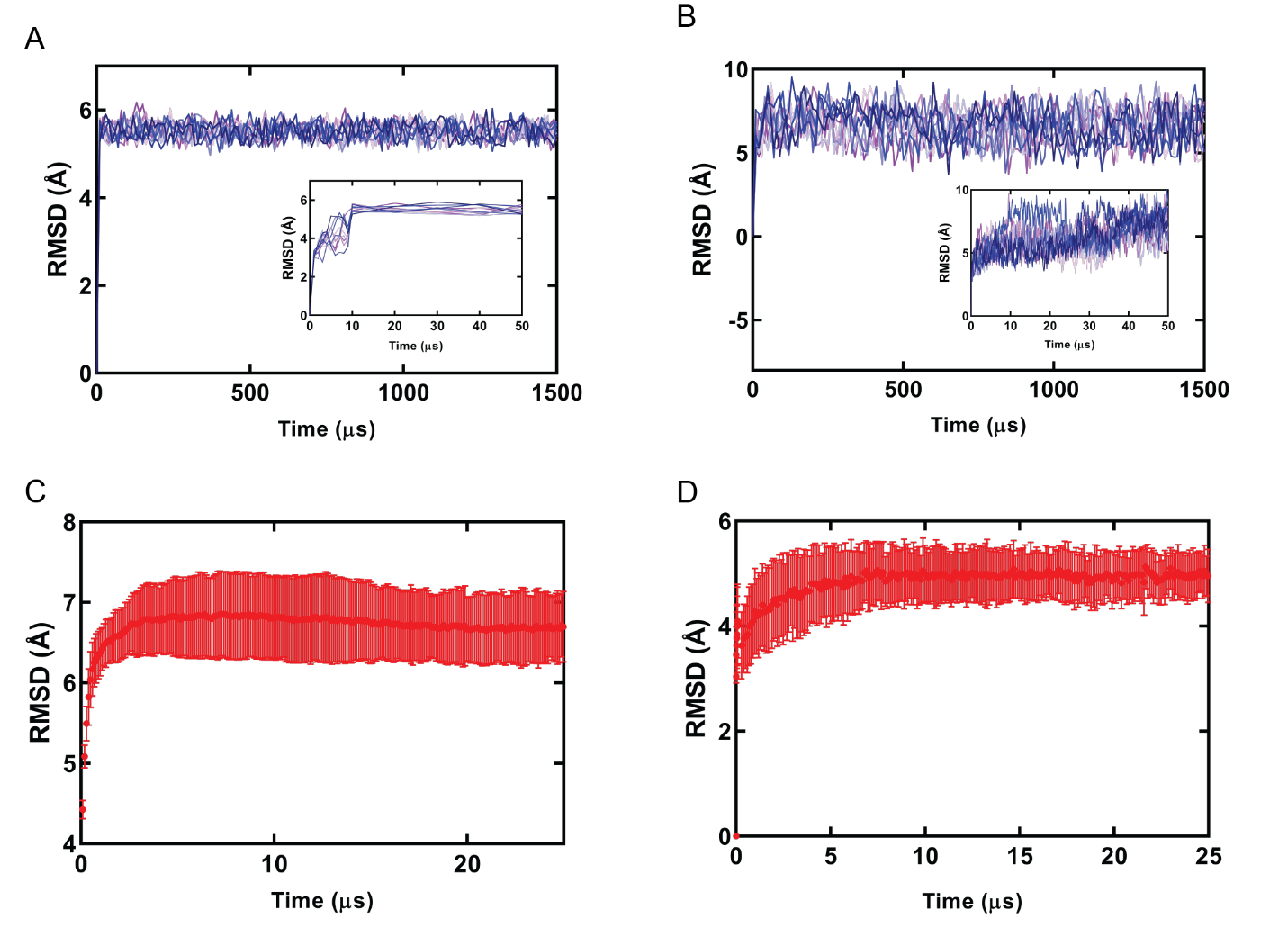

**Supplementary Figure 2 RMSD Convergence of Structure-based simulations**. (A) RMSD of ribosome backbone using potential 1 for 10 different 1.5 ms simulation. Inlet represents the first 50 µs of simulation where the simulation converges at ~10 µs. (B) RMSD of aa-tRNA backbone in simulations using potential 1 for 1.5 ms simulations. Inlet represents the first 50 50 µs of simulation where the simulation converges at ~50 µs. (C) RMSD of ribosome backbone using potential 2 for the 100 simulations at 25 µs, convergence is reached at ~5 µs. (D) RMSD of aa-tRNA backbone using potential 2 for 100 simulations at 25 µs, convergence is reached at ~5 µs.

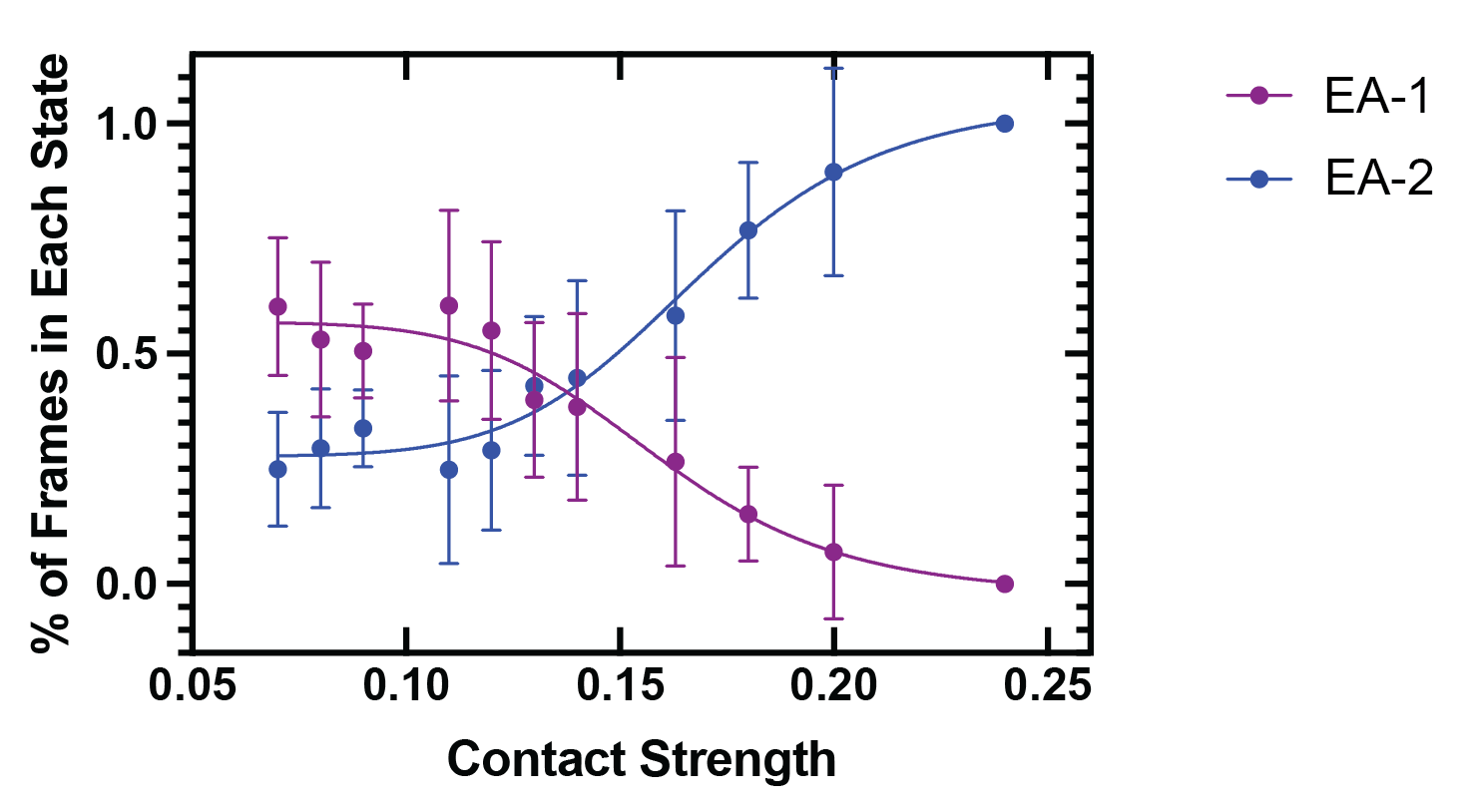

**Supplementary Figure 3 Scaling of A/A native contacts.** Percentage of frames that the simulation was identified to be in the EA-1 or EA-2 position in simulations using potential 1 as measured by the R_elbow_ distance. At a contact weighting of 0.13 the simulations acheived an even distribution of time spent in the EA-1 and EA-2 positions.

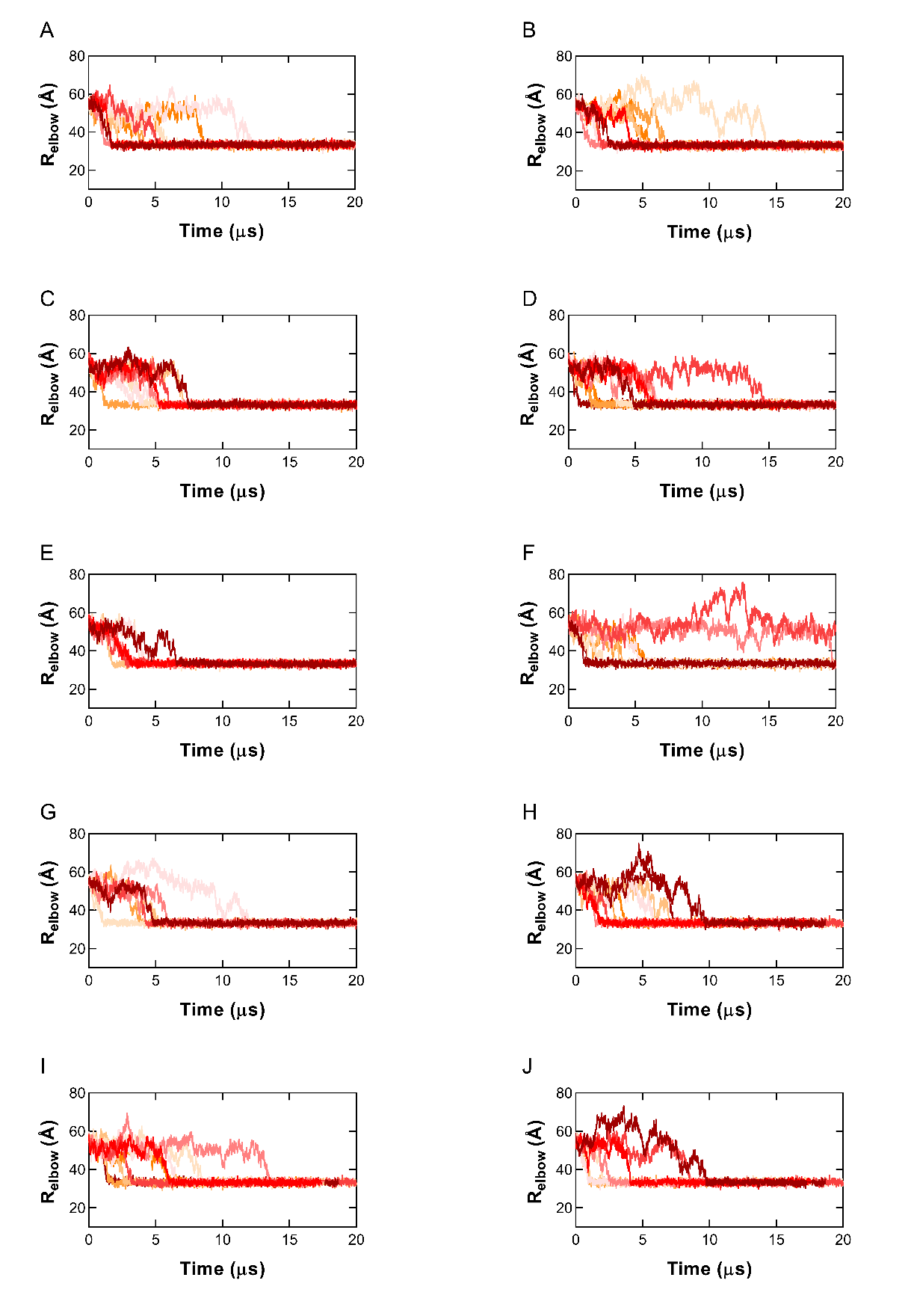

**Supplemental Figure 4 R_elbow_ measurements of 100 Accommodation simulations using potential 2.** R_elbow_ measurements between accommodating aa-tRNA and peptidyl-tRNA during structure-based simulations using potential 2 for simulations 1-10 (A), 11-20 (B), 21-30 (C), 31-40 (D), 41-50 (E), 51-60 (F), 61-70(G), 71-80(H), 81-90(I), 91-100(J).

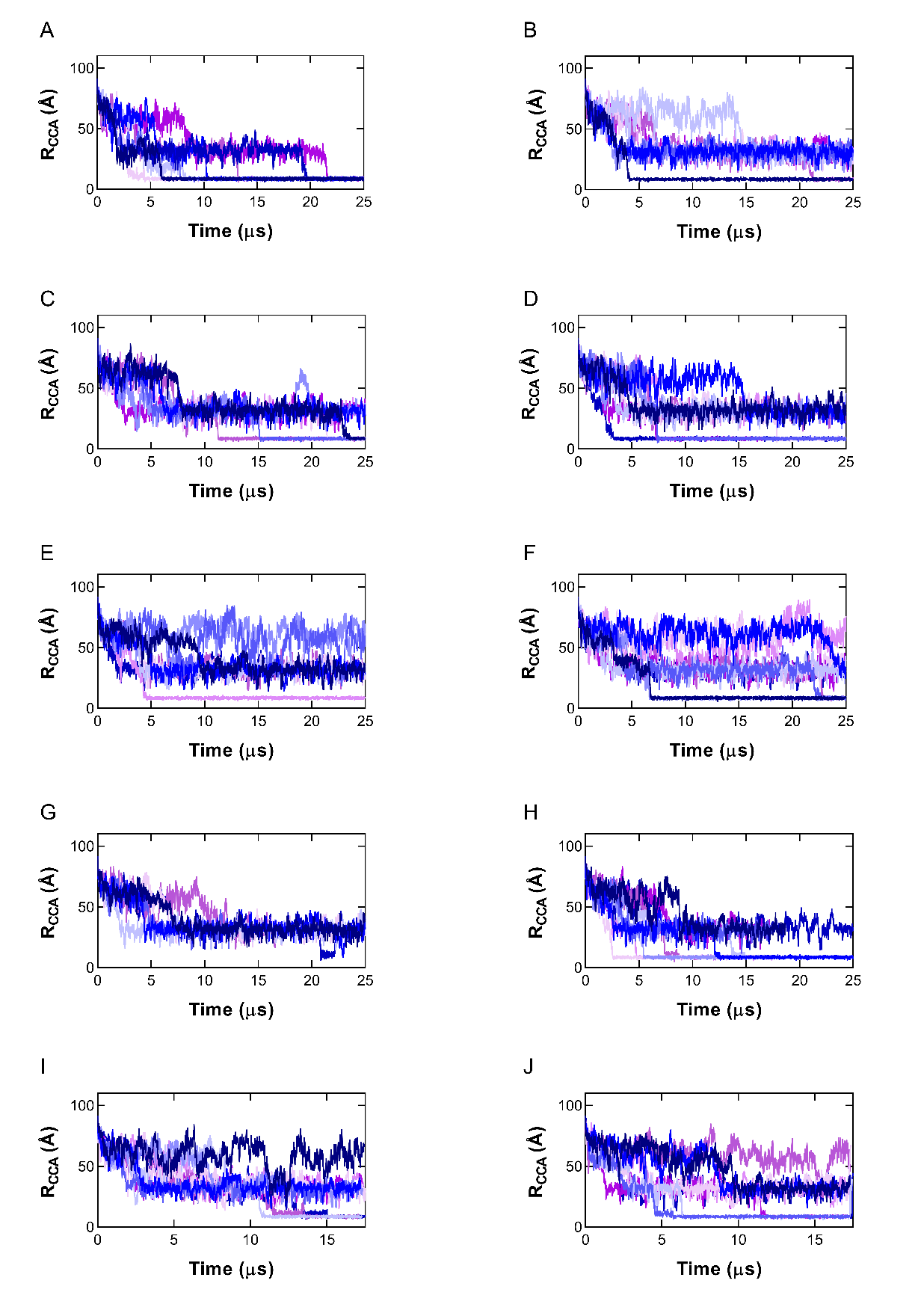

**Supplemental Figure 5 R_CCA_ measurements of 100 Accommodation simulations using potential 2.** R_CCA_ measurements between accommodating aa-tRNA and peptidyl-tRNA during structure-based simulations using potential 2 for simulations 1-10 (A), 11-20 (B), 21-30 (C), 31-40 (D), 41-50 (E), 51-60 (F), 61-70(G), 71-80(H), 81-90(I), 91-100(J).

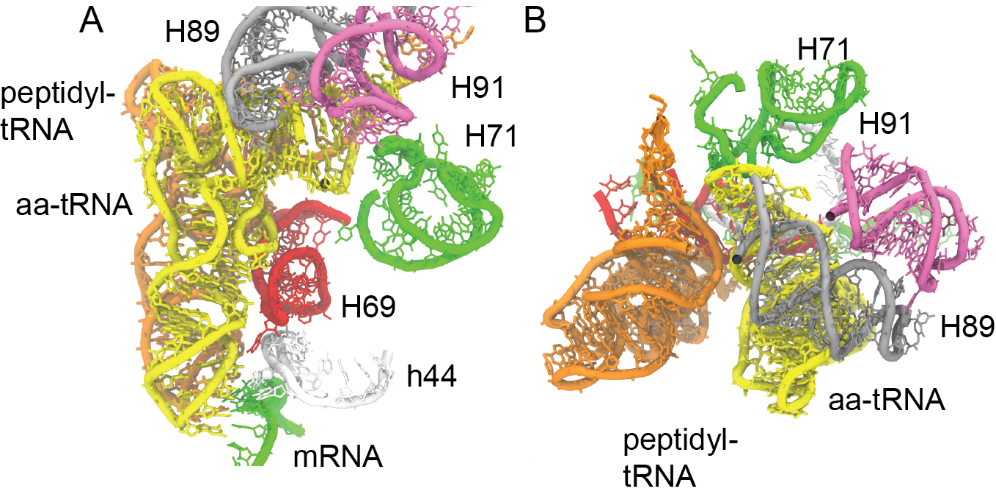

**Supplementary Figure 6 Structural Representation of the R_cca_ intermediate conformation.** (A) All-atom structural representation of aa-tRNA accommodating into the A-site in a position where R_cca_ is in the intermediate position of ~25 Å. View is from the entrance of the A-site. (B) All-atom structural representation of aa-tRNA accommodating into the A-site in a position where R_cca_ is in the intermediate position of ~25 Å. View is from the LSU. In these models aa-tRNA (yellow), peptidyl-tRNA (orange), H71 (green), H69, (red), h44 (white), H89 (grey), H90 (pink), and mRNA (green) are represented.

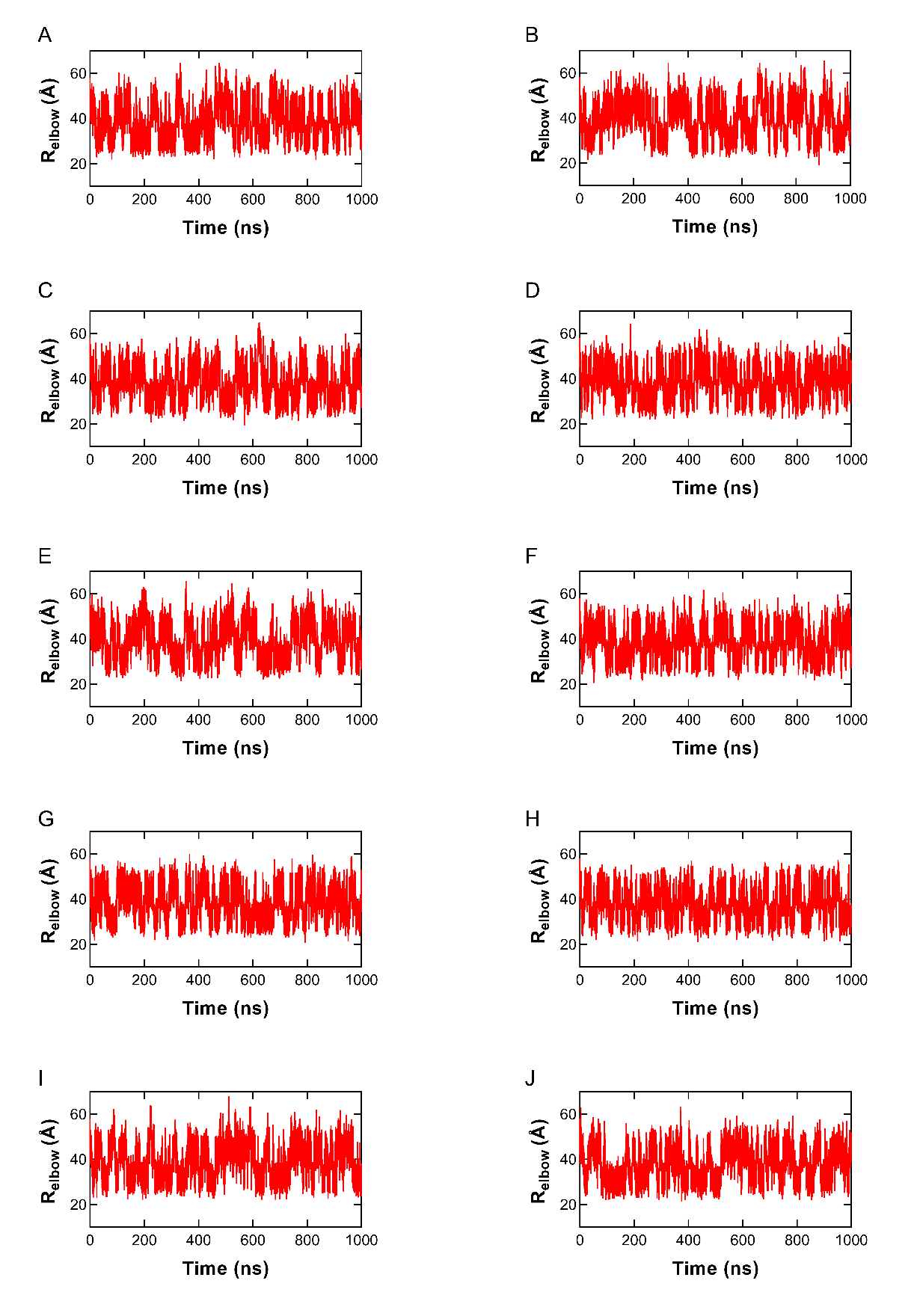

**Supplemental Figure 7 R_elbow_ measurements of 10 Accommodation simulations using potential 1.** R_elbow_ measurements between accommodating aa-tRNA and peptidyl-tRNA during structure-based simulations using potential 1 for simulations 1 (A), 2 (B), 3 (C), 4 (D), 5 (E), 6 (F), 7(G), 8(H), 9(I), 10(J).

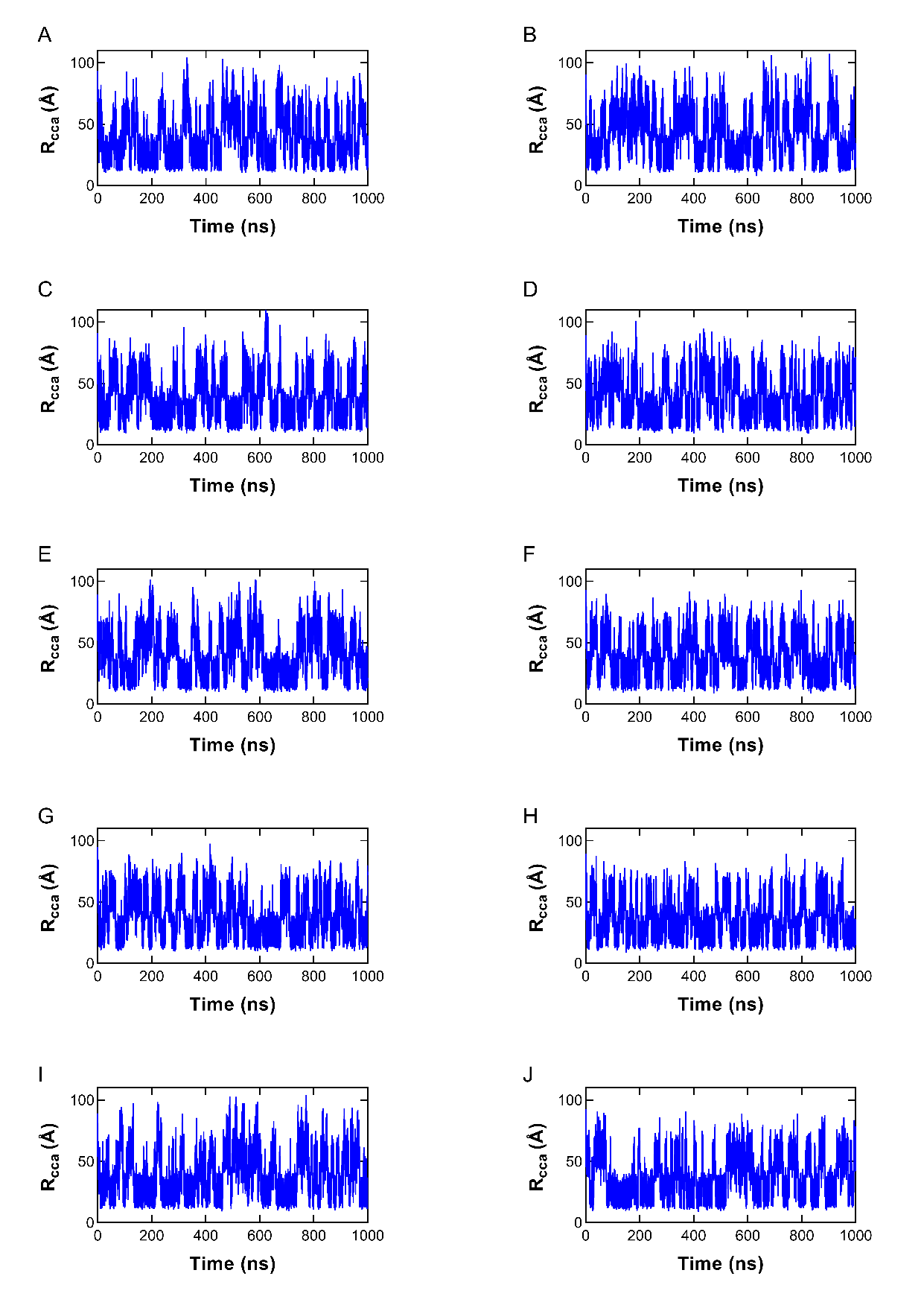

**Supplemental Figure 8 R_CCA_ measurements of 10 Accommodation simulations using potential 1.** R_CCA_ measurements between accommodating aa-tRNA and peptidyl-tRNA during structure-based simulations using potential 1 for simulations 1 (A), 2 (B), 3 (C), 4 (D), 5 (E), 6 (F), 7(G), 8(H), 9(I), 10(J).

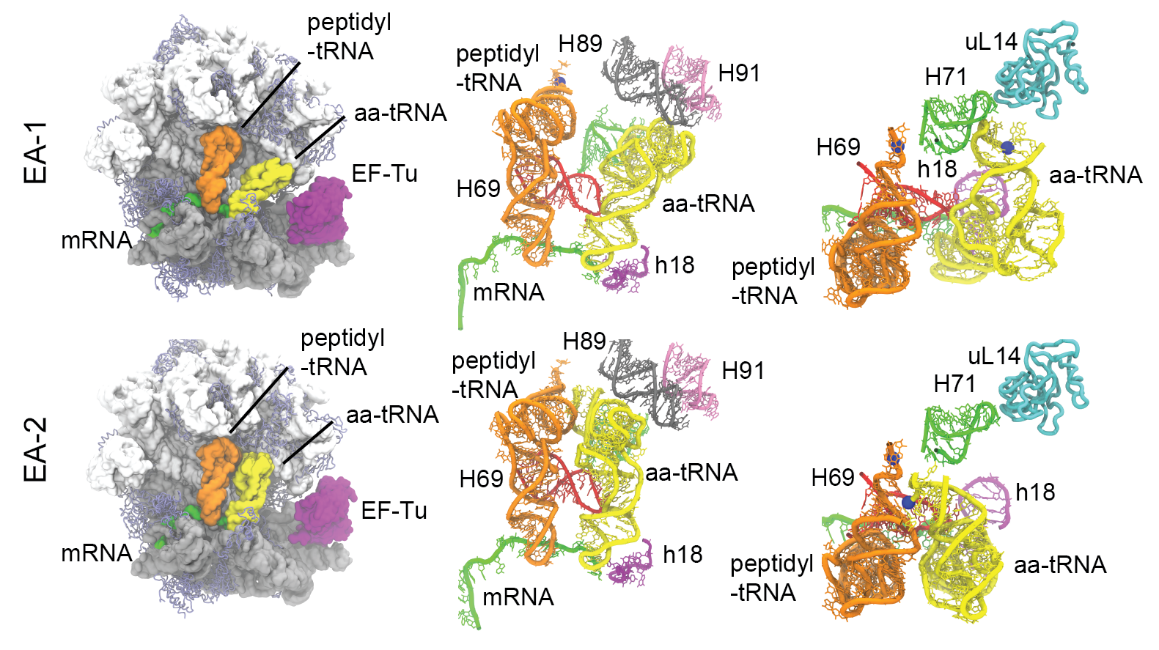

**Supplemental Figure 9. Elbow Accommodated positions observed in *E. coli* accommodation simulations.** Representative structure of EA-1 (top). Complete 70S structure of EA-1 (left). Representation of the accommodation corridor of the ribosome with aa-tRNA engaging H89 in EA-1 (middle). Representation of the accommodation corridor in EA-1 from the LSU perspective (right). Representative structure of EA-2 (bottom). Complete 70S structure of EA-2 (left). Representation of the accommodation corridor of the ribosome with aa-tRNA engaging H71 after passing H89 (middle). Representation of the accommodation corridor in EA-2 from the LSU perspective (right). In these models the rRNA (white and grey), ribosomal proteins (blue), peptidyl-tRNA (orange), aa-tRNA (yellow), H89 (grey), H90 (mauve), H71 (green), h18 (purple), H44 (red), and mRNA (green) are represented.

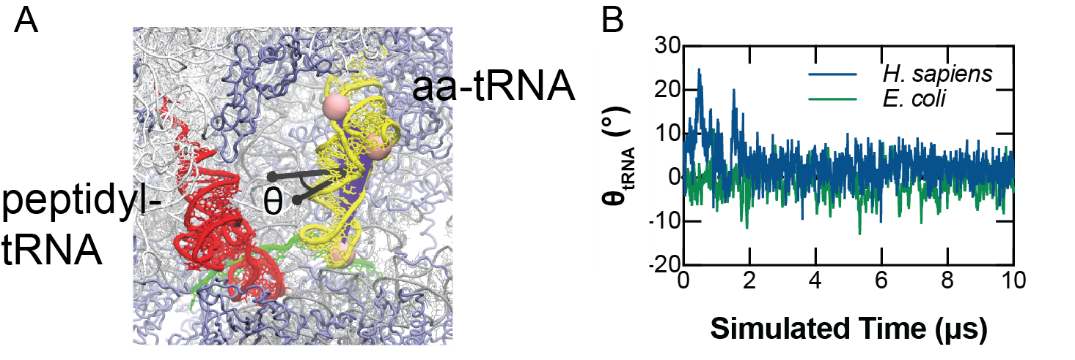

**Supplemental Figure 10. θ_tRNA_ angle measurement.** (A) aa-tRNA pivoting measured by the angle change of the vector (θ) perpendicular to the plane (blue) defined by atoms C4, A35, and G56 (pink) of the accommodating tRNA (yellow). (B) Change in tRNA Angle (θ_tRNA_) of *H. sapiens* and *E. coli* aa-tRNA during accommodation into the ribosome using potential 2.

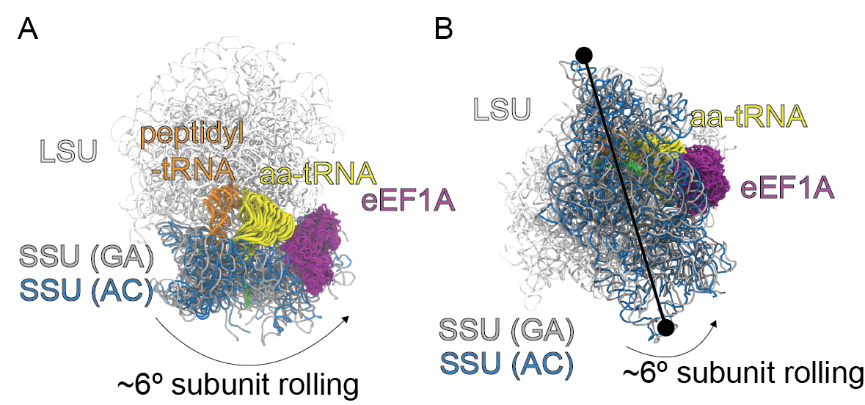

**Supplemental Figure 11. ribosomal SSU rolling.** (A) representative structure of subunit rolling during aa-tRNA accommodation. LSU (white), eEF1A (purple), aa-tRNA (yellow), peptidyl-tRNA (orange), and SSU (GA-silver, AC-blue) are represented. Aa-tRNA is represented in multiple positions during accommodation into the A site. (B) View of the ribosome during subunit rolling from the SSU to highlight the axis of rolling.

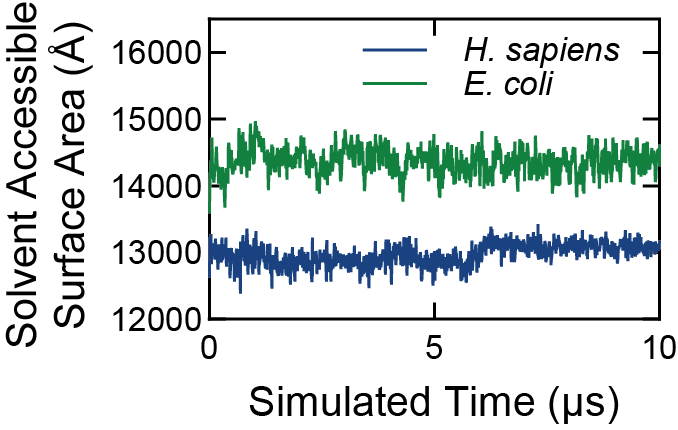

**Supplementary Figure 12. Solvent Accessible Surface Area of aa-tRNA**. Time-dependence of changes in Solvent Accessible Surface Area (SASA) of aa-tRNA during accommodation into the A site of the ribosome. *H. sapiens* and *E. coli* tRNA accommodation represented, indicating that *H. sapiens* have less SASA and that the SASA remains constant during the entirety of the simulation. Data generated from simulations using potential 2.

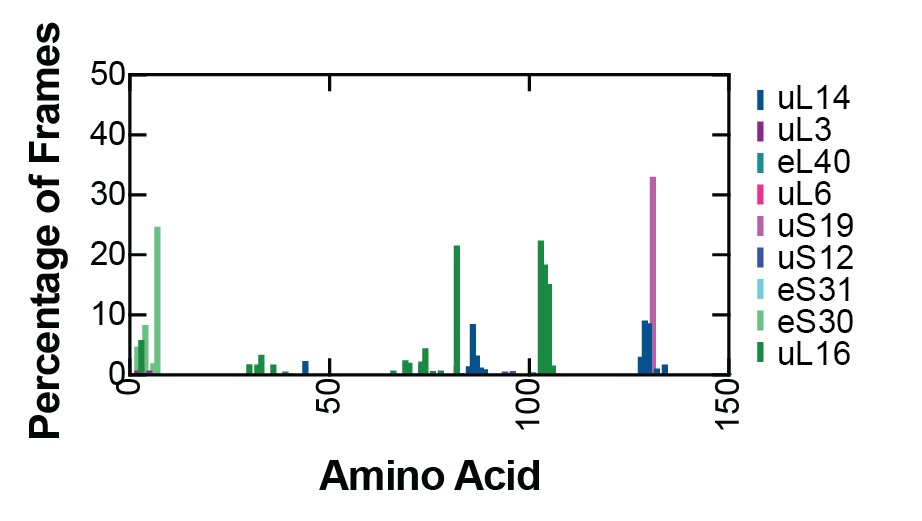

**Supplementary Figure 13. Ribosomal Proteins that are proximal to the accommodating aa-tRNA**. Amino acids that are within 4 Å of the accommodating tRNA during accommodation in the ribosomal accommodation corridor. The percentage of frames that they are within this distance cut off are reported.

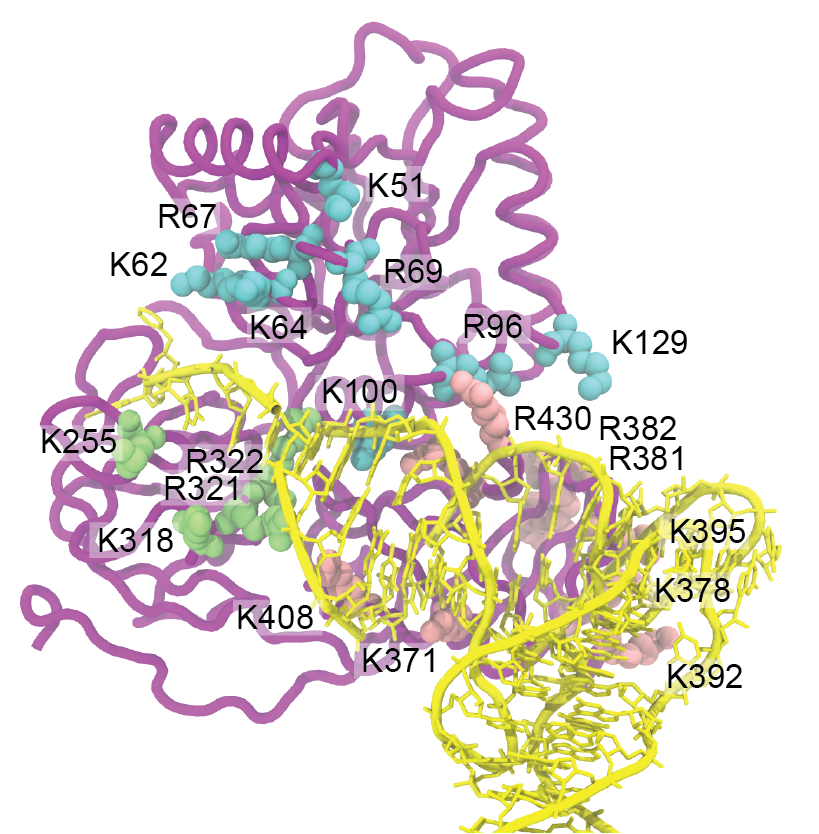

**Supplementary Figure 14. Basic amino acids of eEF1A that are within 4 Å of accommodating aa-tRNA during structure-based simulations.** Basic amino acids that are within the cutoff distance of 4 Å of the accommodating aa-tRNA are highlighted as cyan (Domain I), green (Domain II), or pink (Domain III). Basic amino acids that are in contact with the aa-tRNA in the starting GTPase activated conformation are identified in addition to those in switch I, which interacts with the 3’CCA minor groove, and positive amino acids on the distal face of domain III.

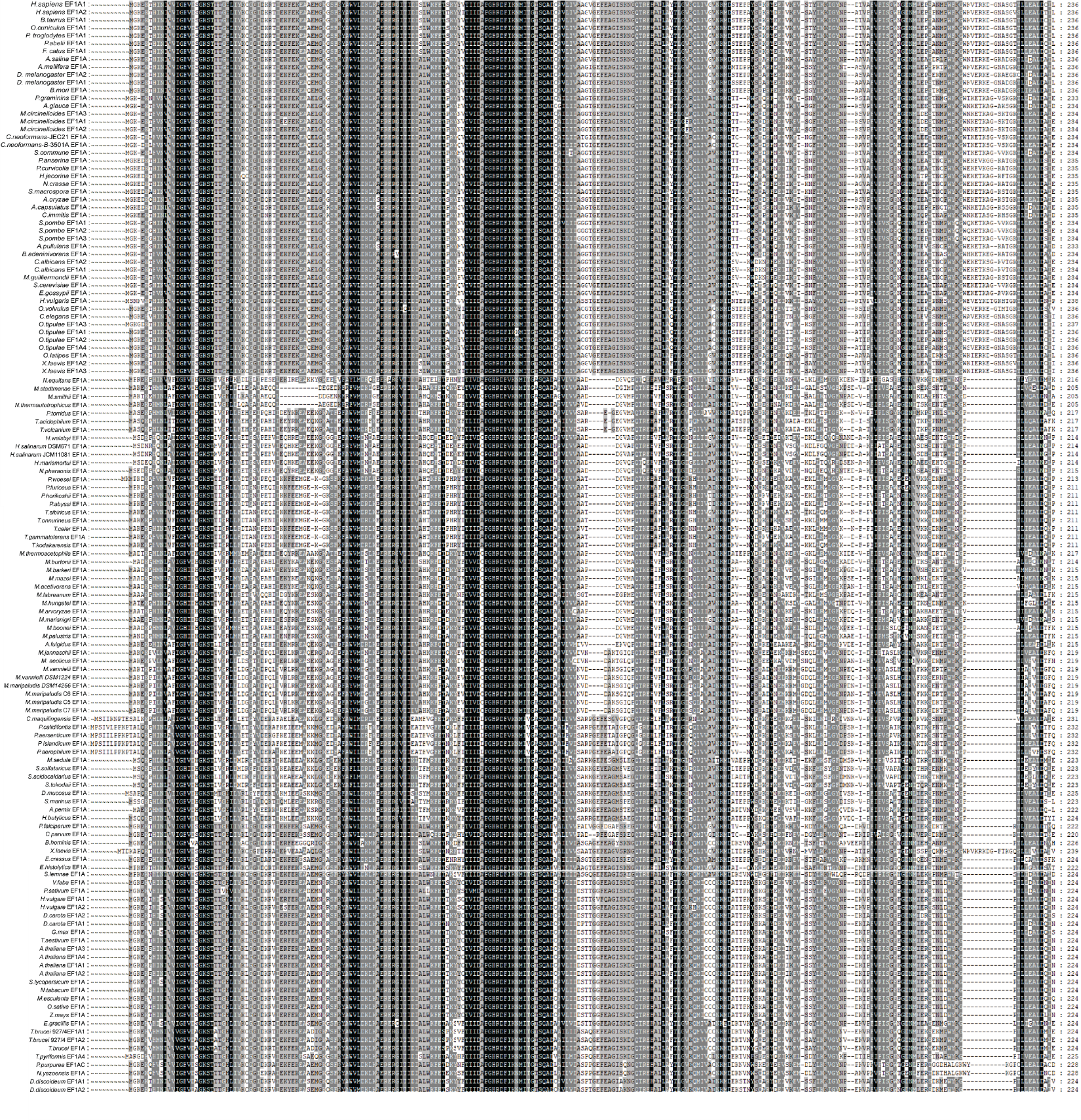

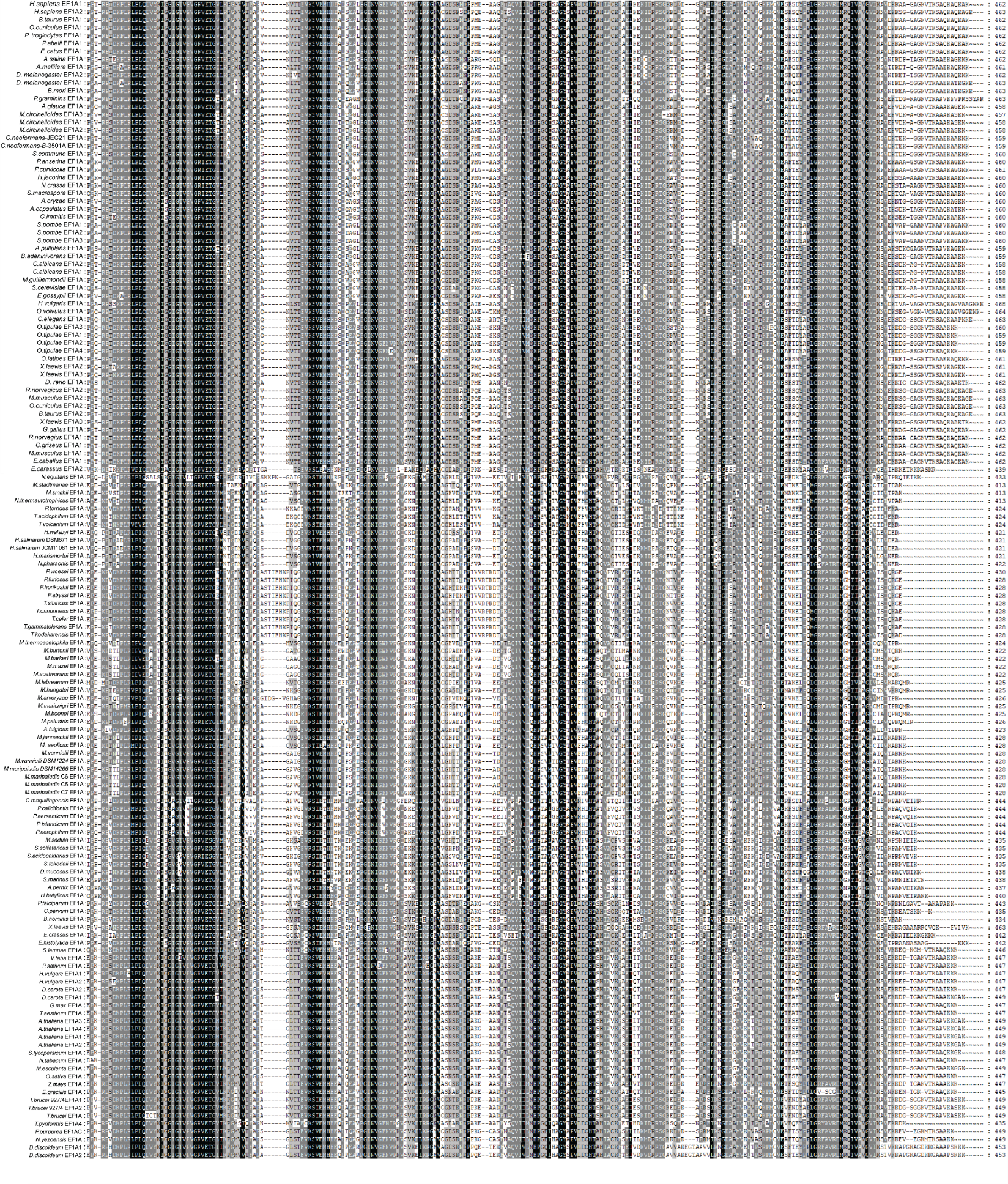

**Supplemental Figure 15 multiple sequence alignment of 147 eEF1A genes from eukaryotic species.** Multiple sequence alignment of eEF1A genes accessed from Uniprot^4^ and aligned using Clustal Omega^5^. Alignment was visualized in Genedoc where amino acids highlighted in black are conserved, those highlighted in grey have similar chemistry and no highlighting indicates no conservation.

**Supplemental Table 1 Barrier estimation of human and *E. coli* aa-tRNA accommodation**. Rates (k) were measured in Holm *et al*. Nature 2023^6^.

|  | A (x 10^-6^ s^-1^) | k at 25 °C^6^ (s^-1^) | E_A_ (K_B_T) |
| --- | --- | --- | --- |
| *H. sapiens* | 5.1 ± 4.0 | 1.7 ± 0.2 | 11-13.4 |
| *E. coli* | 2.1 ± 1.1 | 30 | 9.3-10.4 |

**Supplemental Table 2. Conservation of tRNA Nucleotides that interact with Domain III of eEF1A.** tRNA sequences from *H. sapiens* (GRCh37/hg19) are compared to determine conservation within humans. tRNA sequences from various species including *H. sapiens* (GRCh37/hg19), *S. cerevisiae* (S288C), *M. musculus* (GRCm39/mm39), *B. subtillus* (subsp subtilis str 168), *E. coli* (str k-12 substr. MG1655), *D. melanogaster* (BDGP Rel. 6/dm6), *A. thaliana* (TAIR10), *S. pombe* (972h-) are compared to determine conservation across domains of life. All tRNA sequences were accessed from the GtRNA database^7^ aligned in Clustal Omega^8^ and analyzed with Genedoc.

|  | *H. sapiens* tRNA | Various species tRNA |
| --- | --- | --- |
| G1 | 84.69% | 80.89% |
| U51 | 37.5% | 66.98% |
| G52 | 6.45% | 0.43% |
| A64 | 10.44% | 24.15% |
| G65 | 99.31% | 47.37% |
| U66 | 92.13% | 31.48% |

**Supplemental Table 3. Information for Structure-based simulations.** Information for structure-based simulations. Water atoms were implicitly defined, and no salt was added as electrostatic contributions were not considered in the model.

|  | Box Dimensions (Å^3^) | Number of Atoms | Added Salt (M) | Number of Timesteps | Estimated Simulated Time (ms) | Number of Simulations |
| --- | --- | --- | --- | --- | --- | --- |
| Human Potential 1 | 1.25 x 10^5^ | 214569 | 0 | 5 - 7.5 x 10^8^ | 1000-1500 | 10 |
| Human Potential 2 | 1.25 x 10^5^ | 214497 | 0 | 8.5-12.5 x 10^6^ | 17-25 | 100 |
| E. coli Potential 2 | 1.25 x 10^5^ | 154401 | 0 | 12.5 x 10^7^ | 25 | 10 |

1	Whitford, P. C. *et al.* Accommodation of aminoacyl-tRNA into the ribosome involves reversible excursions along multiple pathways. *RNA* **16**, 1196-1204 (2010). <https://doi.org/10.1261/rna.2035410>

2	Geggier, P. *et al.* Conformational sampling of aminoacyl-tRNA during selection on the bacterial ribosome. *J Mol Biol* **399**, 576-595 (2010). <https://doi.org/10.1016/j.jmb.2010.04.038>

3	Girodat, D., Wieden, H. J., Blanchard, S. C. & Sanbonmatsu, K. Y. Geometric alignment of aminoacyl-tRNA relative to catalytic centers of the ribosome underpins accurate mRNA decoding. *Nat Commun* **14**, 5582 (2023). <https://doi.org/10.1038/s41467-023-40404-9>

4	UniProt, C. UniProt: the Universal Protein Knowledgebase in 2025. *Nucleic Acids Res* **53**, D609-D617 (2025). <https://doi.org/10.1093/nar/gkae1010>

5	Madeira, F. *et al.* The EMBL-EBI Job Dispatcher sequence analysis tools framework in 2024. *Nucleic Acids Res* **52**, W521-W525 (2024). <https://doi.org/10.1093/nar/gkae241>

6	Holm, M. *et al.* mRNA decoding in human is kinetically and structurally distinct from bacteria. *Nature* **617**, 200-207 (2023). <https://doi.org/10.1038/s41586-023-05908-w>

7	Chan, P. P. & Lowe, T. M. GtRNAdb 2.0: an expanded database of transfer RNA genes identified in complete and draft genomes. *Nucleic Acids Res* **44**, D184-189 (2016). <https://doi.org/10.1093/nar/gkv1309>

8	Sievers, F. *et al.* Fast, scalable generation of high-quality protein multiple sequence alignments using Clustal Omega. *Mol Syst Biol* **7**, 539 (2011). <https://doi.org/10.1038/msb.2011.75>
